# Inhibitory microcircuits for top-down plasticity of sensory representations

**DOI:** 10.1101/494989

**Authors:** Katharina Anna Wilmes, Claudia Clopath

## Abstract

Humans and animals are remarkable at detecting stimuli that predict rewards. While the underlying neural mechanisms are unknown, reward influences plasticity of sensory representations in early sensory areas. The underlying changes in excitatory and inhibitory circuitry are however unclear. Recently, experimental findings suggest that the inhibitory circuits can regulate learning. In addition, the inhibitory neurons in superficial layers are highly modulated by diverse long-range inputs, including reward signals. We, therefore, hypothesise that plasticity of interneuron circuits plays a major role in adjusting stimulus representations. We investigate how top-down modulation by rewards can interact with local excitatory and inhibitory plasticity to induce long-lasting changes in sensory circuitry. Using a computational model of layer 2/3 primary visual cortex, we demonstrate how interneuron networks can store information about the rewarded stimulus to instruct long-term changes in excitatory connectivity in the absence of further reward. In our model, stimulus-tuned somatostatin-positive interneurons (SSTs) develop strong connections to parvalbumin-positive interneurons (PVs) during reward presentation such that they selectively dis-inhibit the pyramidal layer henceforth. This triggers plasticity in the excitatory neurons, which leads to increased stimulus representation. We make specific testable predictions in terms of the activity of different neuron types. We finally show that this two-stage model allows for translation invariance of the learned representation.

## Introduction

Animals learn better when it matters to them. For example, they learn to discriminate sensory stimuli when they receive a reward. As a result of learning, neural responses to sensory stimuli are adjusted even in primary sensory areas, such as primary visual cortex (V1, Goltstein et al., 2013; Khan et al., 2018; Poort et al., 2015). When mice consistently receive a reward after seeing a grating of a given orientation, the tuning preference of layer 2/3 neurons for this rewarded orientation is increased (Goltstein et al., 2013; Poort et al., 2015). It is thought that behaviourally relevant contexts, such as rewards, trigger an internal top-down signal available to these early sensory circuits. This could be mediated by cholinergic inputs from the basal forebrain, for example (Chubykin et al., 2013; Letzkus et al., 2011). By top-down signal, we mean any long-range input to the superficial layers that delivers behaviorally relevant information to the local circuit.

Pyramidal cells in primary sensory cortices are embedded in a canonical microcircuit motif with different types of inhibitory interneurons. The main inhibitory types are the PVs, the SSTs, and the VIPs. Top-down inputs project to superficial layers (Atallah et al., 2012; Fu et al., 2014; Lee et al., 2013; Petro et al., 2014; Pi et al., 2013; Zhang et al., 2014). They target multiple cell types. For example, VIPs in the primary auditory cortex are activated when a reward is present (Pi et al., 2013). Inhibitory synapses are plastic (see (Vogels et al., 2013) for a review) and perturbation of interneurons impairs learning (Letzkus et al. (2011), see (Lucas and Clem, 2018) for a recent review).

We hypothesised that the inhibitory circuitry in layer 2/3 mediates the top-down instructions (e.g triggered by a reward) to guide slow plastic changes in the circuit beyond the presence of reward. We wanted to test whether interneurons can learn from a top-down signal. The inhibitory connectivity structure could then instruct the excitatory cells in the absence of top-down modulation. To test this, we built a biologically constrained computational model of layer 2/3 primary visual cortex. We simulated a rewarded phase in which the presentation of one stimulus is paired with a reward signal, which excites VIPs. We then simulated a second refinement phase, where the sensory stimuli were presented without the reward. During the first rewarded phase, connections between SSTs and PVs developed a specific connectivity structure. This structure triggered disinhibition of the excitatory neurons even in the absence of reward. Plasticity in the excitatory neurons, therefore, shaped the microcircuit during the second refinement phase. It led to an increased stimulus preference of the previously rewarded stimulus. Our model offers testable predictions on the activity of different cell types during and after the reward presentation. We also propose that this two-stage mechanism allows for learned representations to generalise across different parts of the visual space.

## Results

### Hypothesis: Two-stage model of top-down guided microcircuit plasticity

Neural responses to visual stimuli in V1 are not a simple function of bottom-up sensory inputs. They are additionally modulated by various inputs from other areas (Khan and Hofer, 2018; Pakan et al., 2018) and by recurrent local excitatory and inhibitory neurons (especially in layer 2/3, Cossell et al., 2015). We hypothesised that top-down inputs can induce changes in sensory representations via changes in recurrent connections in two stages.

i. *The rewarded phase*. A specific stimulus (e.g. a vertical bar) is paired with a reward-mediated top-down signal which excites VIPs (triple arrow in Fig. 1a top). The VIPs inhibit the SSTs, which we assume are stimulus-tuned (Cottam et al., 2013; Ma et al., 2010). At the same time, the VIPs disinhibit the PVs, which we model as untuned (Cottam et al., 2013; Ma et al., 2010). Activity-dependent plasticity then increases the connections between SSTs which are tuned to the rewarded stimulus (here the vertical bar, vertSSTs) and PVs (Fig. 1b top). The inhibitory motif now carries information about the reward. In addition, this inhibitory structure disinhibits the excitatory neurons (Fig. 1b bottom).

**Figure 1:**
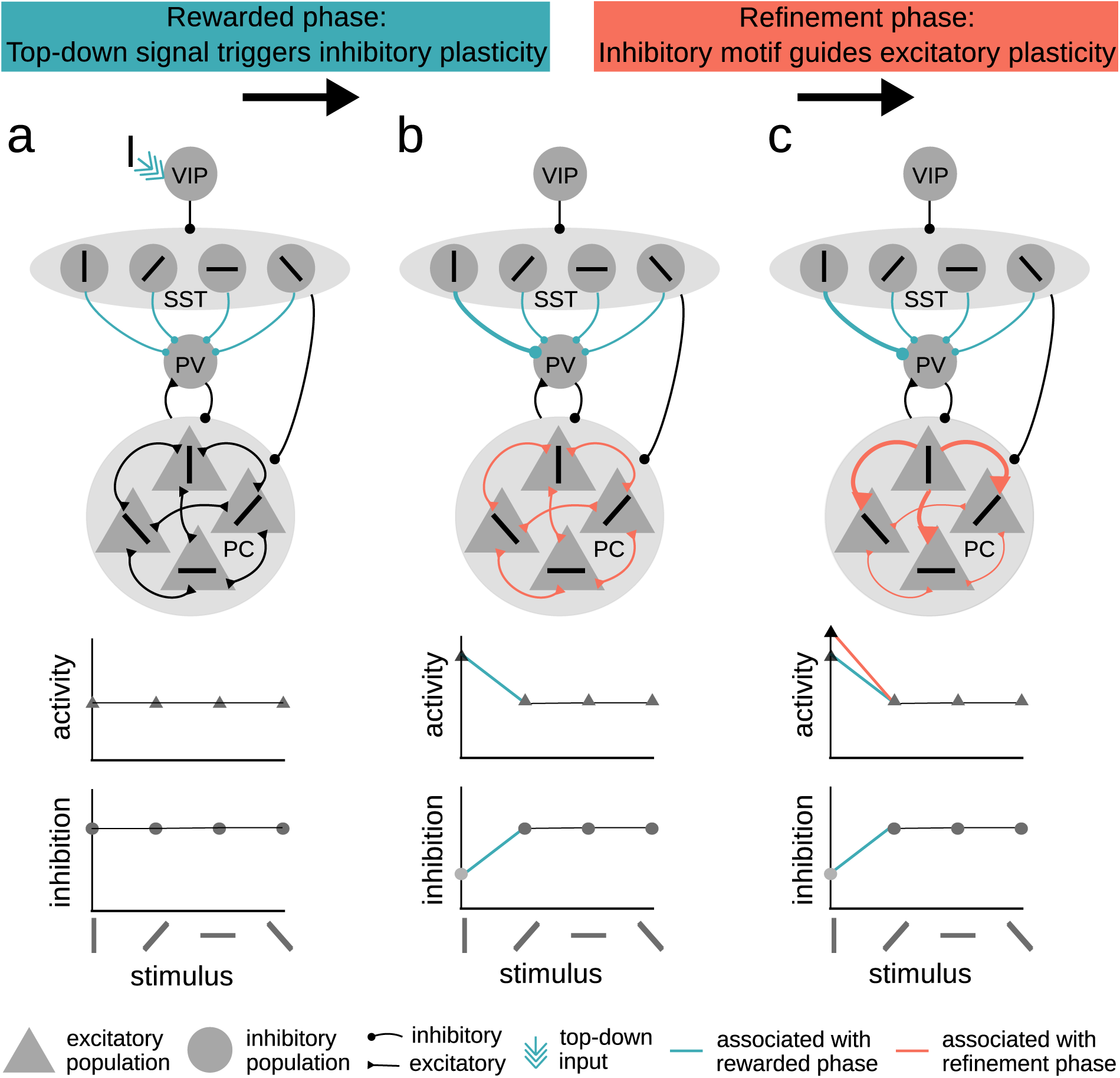
The two-stage model of top-down guided plasticity. a: Before the rewarded phase. We assume the SSTs and PCs to be stimulus-tuned and PVs to be untuned. During the rewarded phase, the top-down signal activates VIPs when the rewarded stimulus is present (vertical bar). This triggers plasticity at the SST-to-PV connections. b: At the end of the rewarded phase and at the beginning of the refinement phase, there are strong connections from the SSTs tuned to the vertical bar to PVs (green, top). The PV activity is therefore low for the vertical bar (bottom). The excitatory neurons coding for the vertical bar are disinhibited (middle). During the refinement phase, the inhibitory motif guides plasticity at the excitatory neurons. c: At the end of the refinement phase, strong connections from the excitatory neurons coding for the vertical bar to the other excitatory neurons have developed (red, top). This results in an increased activity of excitatory neurons towards the vertical bar (red line, middle) beyond that resulting from reduced inhibitory PV activity (blue line, middle, and bottom).
ii. *The refinement phase*. In the second phase, the reward and therefore also the top-down input is absent (Fig. 1b top). As the inhibitory (SST-PV) motif disinhibits the PCs (Fig. 1b top), it opens a window for plasticity at the excitatory synapses. This will result in a refinement of the excitatory connectivity. Strong recurrent connections from PCs coding for the vertical bar (vertPC) to other excitatory neurons will develop. All PCs will, therefore, have an increased response to the vertical bar stimulus.

In summary, we hypothesised that learning can happen in two stages. To test this, we simulated a mechanistic model of the layer 2/3 microcircuit.

### Top-down signal triggers plasticity in the inhibitory circuit

We simulated a spiking neural network model of the canonical microcircuit of layer 2/3 mouse primary visual cortex (Pfeffer et al. 2013). Neurons were modelled as integrate-and-fire neurons (Pfeffer et al., 2013). VIPs inhibited the SSTs, which in turn inhibited the PVs and the PCs. The PVs inhibited the PCs. The PCs were recurrently connected (Fig. 1) (Pfeffer et al., 2013). PCs and SST were tuned to orientation (Cottam et al., 2013; Ma et al., 2010, but see Kerlin et al. (2010)). Recurrent excitatory connections and those from SSTs to PVs were plastic according to the classical spike-timing-dependent plasticity (STDP) model. All other connections were fixed (see methods for details).

Before we tested our hypothesis, we needed to bring our model from random initial connectivity to a set of weights that corresponds to adult V1 connectivity. We call that the *developmental phase*. During this phase, we randomly presented inputs to our network corresponding to oriented gratings. Excitatory neurons that code for the same orientation were coactive. Therefore, they formed strong clusters due to Hebbian learning (Clopath et al., 2010; Ko et al., 2013) (Fig. 2f developmental phase, Fig. 2e level middle). The SST-to-PV weights did not form a specific structure during the developmental phase (Fig. 2b developmental phase) consistent with experimental literature (Guan et al., 2017).

**Figure 2:**
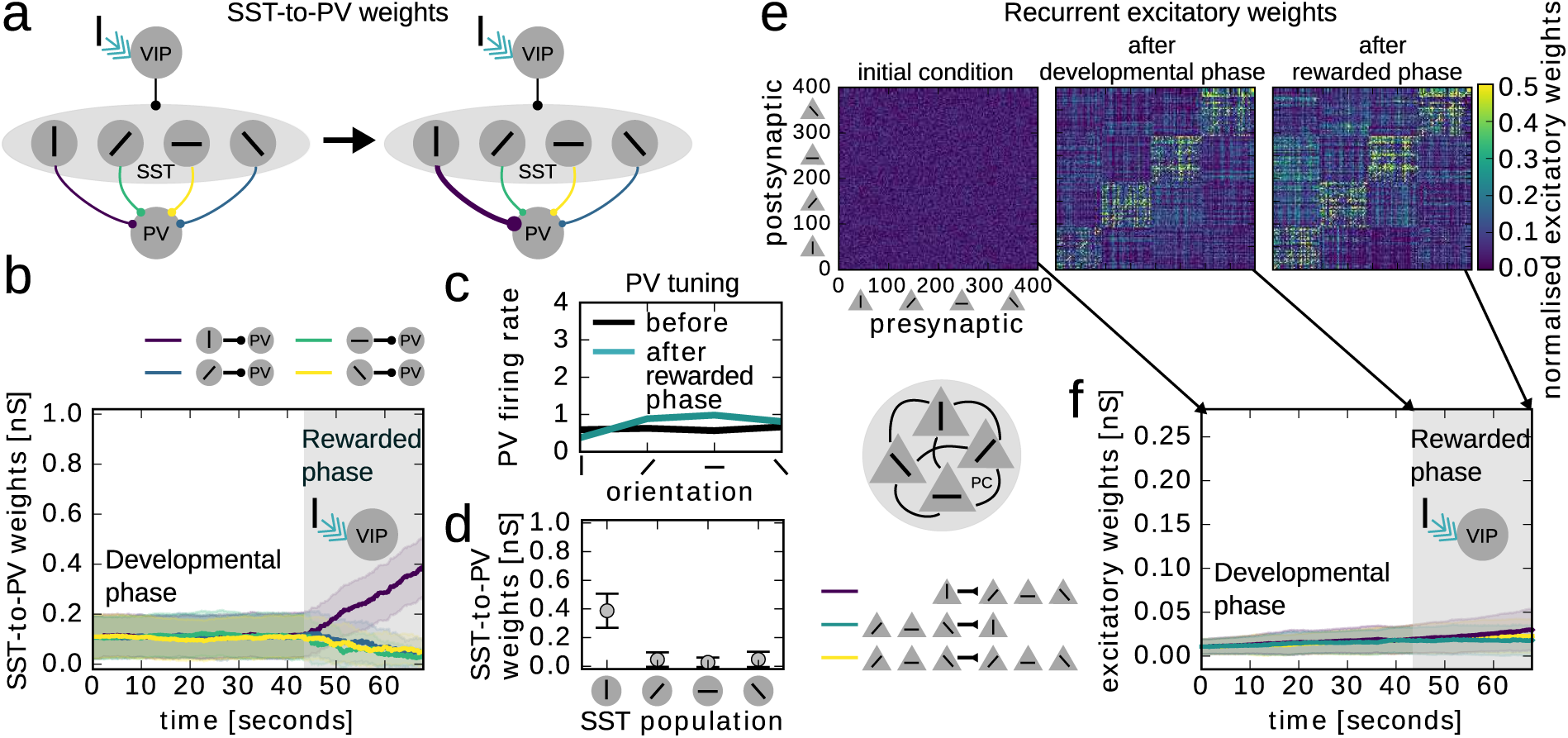
Top-down signal triggers fast plasticity in the inhibitory circuit. a: Illustration of changes in the inhibitory structure during top-down modulation (cyan arrow). b: Evolution of the inhibitory SST-to-PV connections, grouped according to SST tuning preference (colours match the connections in a, shown are the mean and standard deviation). c: Tuning of PVs after the rewarded phase. d: SST-to-PV weights (mean and standard deviation) at the end of the rewarded phase averaged over SST tuning preference. e: Recurrent excitatory weights at different time points in the simulation (initial random condition, after the developmental phase, after the rewarded phase). f: Evolution of excitatory connections (mean and standard deviation). Vertically tuned PCs to vertically tuned PCs (vertPC-vertPC green), vertically tuned PCs to non-vertically tuned PCs (vertPC-nvert-PC, purple), non-vertically tuned PCs to non-vertically tuned PCs (nvertPC-nvert-PC, yellow).

We then simulated a rewarded phase (grey background in Fig. 2b and f). A reward signal excited the VIP population when the vertical bar stimulus was present. This top-down signal was in itself untuned. However, the temporal coincidence with the vertical bar made it stimulus-specific. Connections from the vertical-bar tuned SSTs (vertSST) to PVs increased (purple line in Fig. 2b, rewarded phase). The resulting SST-to-PV structure (Fig. 2d) carried information about the identity of the rewarded stimulus. Hence, the PVs became less responsive to the rewarded stimulus (vertical bar, Fig. 2c). Notably, no significant stimulus-specific structure arose between excitatory connections (Fig. 2e and f). Accordingly, the tuning of excitatory populations did not change.

In summary, unspecific top-down signals can induce an inhibitory connectivity structure without changing excitatory connectivity.

### Inhibitory structure guides excitatory plasticity in the absence of reward

We then tested whether the interneuron structure can guide plasticity in the excitatory neurons in the absence of reward. After the rewarded phase, the inhibitory structure effectively disinhibited all PCs when a vertical bar was present (corresponding to the previously rewarded stimulus). The vertically tuned PCs fired a few milliseconds before the other PCs because they received additional feedforward inputs. STDP, therefore, led to a strengthening of the connections from the vertically tuned PCs to the other PCs (Fig. 3b purple line). Accordingly, connections in the reverse direction were depressed (Fig. 3b green line; see Fig. S1 for spiking details). As a result of the excitatory connectivity structure (Fig. 2c), all PC populations showed an increased response to the vertical bar (Fig. 3d). Since PCs are driving the PVs, PVs became tuned to the vertical bar (Fig. 3h). Synergistically, SST-to-PV connections were strengthened even further (Fig. 3f).

**Figure 3:**
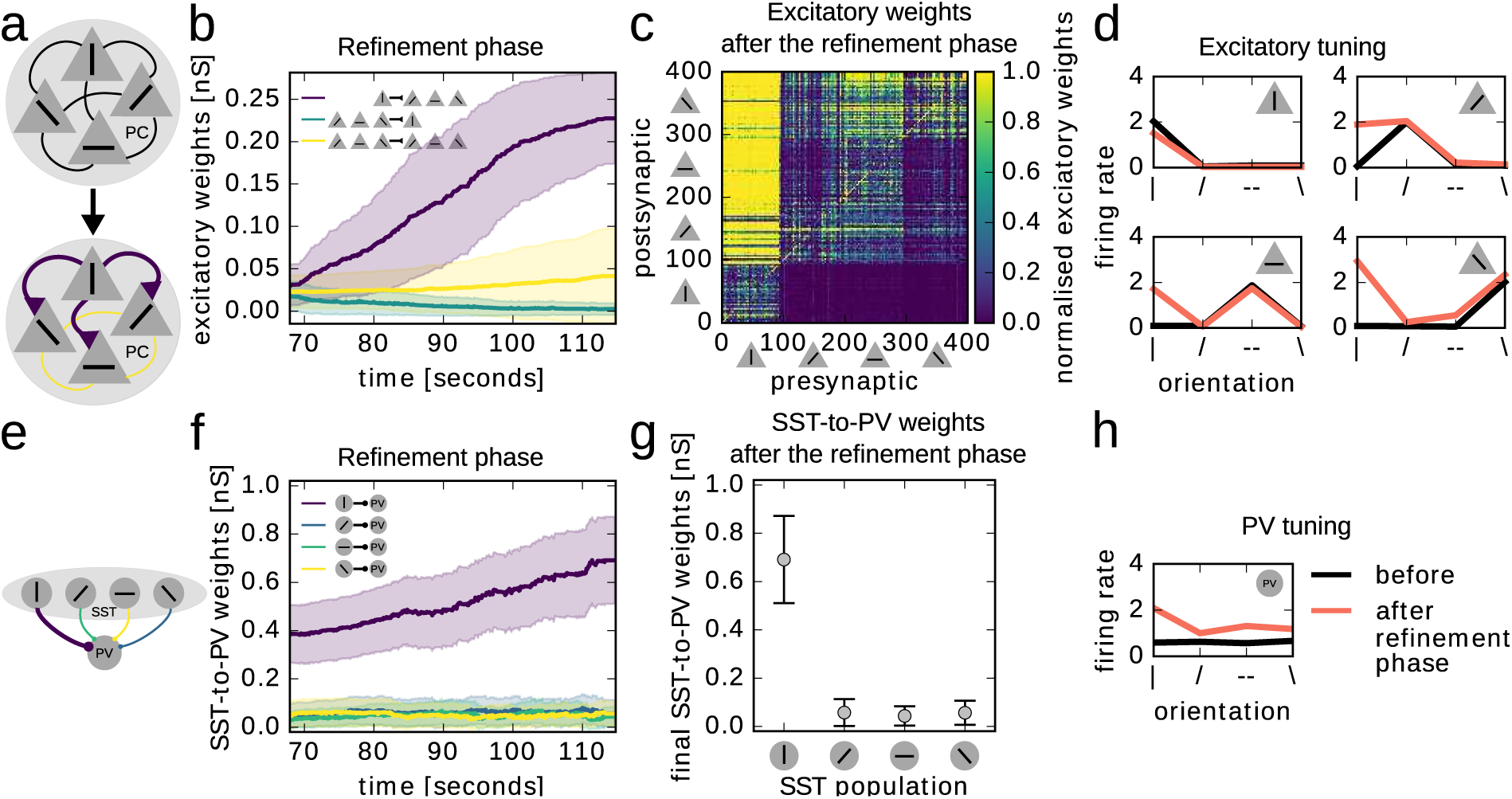
Inhibitory structure guides excitatory plasticity in the absence of reward. a: Illustration of changes in the excitatory structure. b: Evolution of excitatory connections. Mean and standard deviation of the connections from the vertically tuned PCs to PCs tuned to other orientations (vertPC-nvertPC, purple), from PCs tuned to other orientations to the vertically tuned PC population (nvertPC-vertPC, green), and from others to others (nertPC-nvertPC, yellow). c: Final excitatory weight matrix. Neuron 1-100 are tuned to the vertical bar, 100 to 200 to an angled bar, etc. d: Tuning of excitatory populations before the rewarded phase (black) and at the end of the refinement phase (red) (number of spikes during 50 ms after stimulus onset averaged over all occurrences of that stimulus in 1 s of simulation). e: Illustration of the inhibitory structure after the rewarded phase. f: Evolution of the SST-to-PVs connections (mean and standard deviation), grouped according to SST tuning (colours match the colours of the connections in e). g: SST-to-PV connections after the refinement phase, grouped by SST tuning (error bars -standard deviation). h: Tuning of PVs after the refinement phase.

Note, the excitatory structure was stable even if we artificially deleted the inhibitory structure (Fig. S6). The total effect on the PCs depended on the comparative strength of SST-PC and SST-PV-PC pathways. Indeed, we found that the degree to which excitatory structure developed depended on the strength of SST-to-PC connections (Fig. S3f). Finally, we also showed that precise spike timing was not necessary for our two-stage model (see a rate-based implementation in suppl. mat Fig. S7).

In summary, the inhibitory network structure can induce changes in sensory representation by guiding excitatory plasticity.

### Experimentally testable predictions

Our model makes eight precise experimentally testable predictions. (1) PCs and (2) PVs become more tuned to the rewarded stimulus (Fig. 2d, g). (3) SSTs and (4) VIPs do not change their tuning (Fig. S2). (5) Both PC and (6) PV firing rate responses to the rewarded stimulus increase relative to other stimuli (Fig. 4a, b). (7) Excitatory currents increase during the rewarded stimulus (Fig. 4d, S7g), (8) but not the E/I ratio (Fig. 4e, S7i).

**Figure 4:**
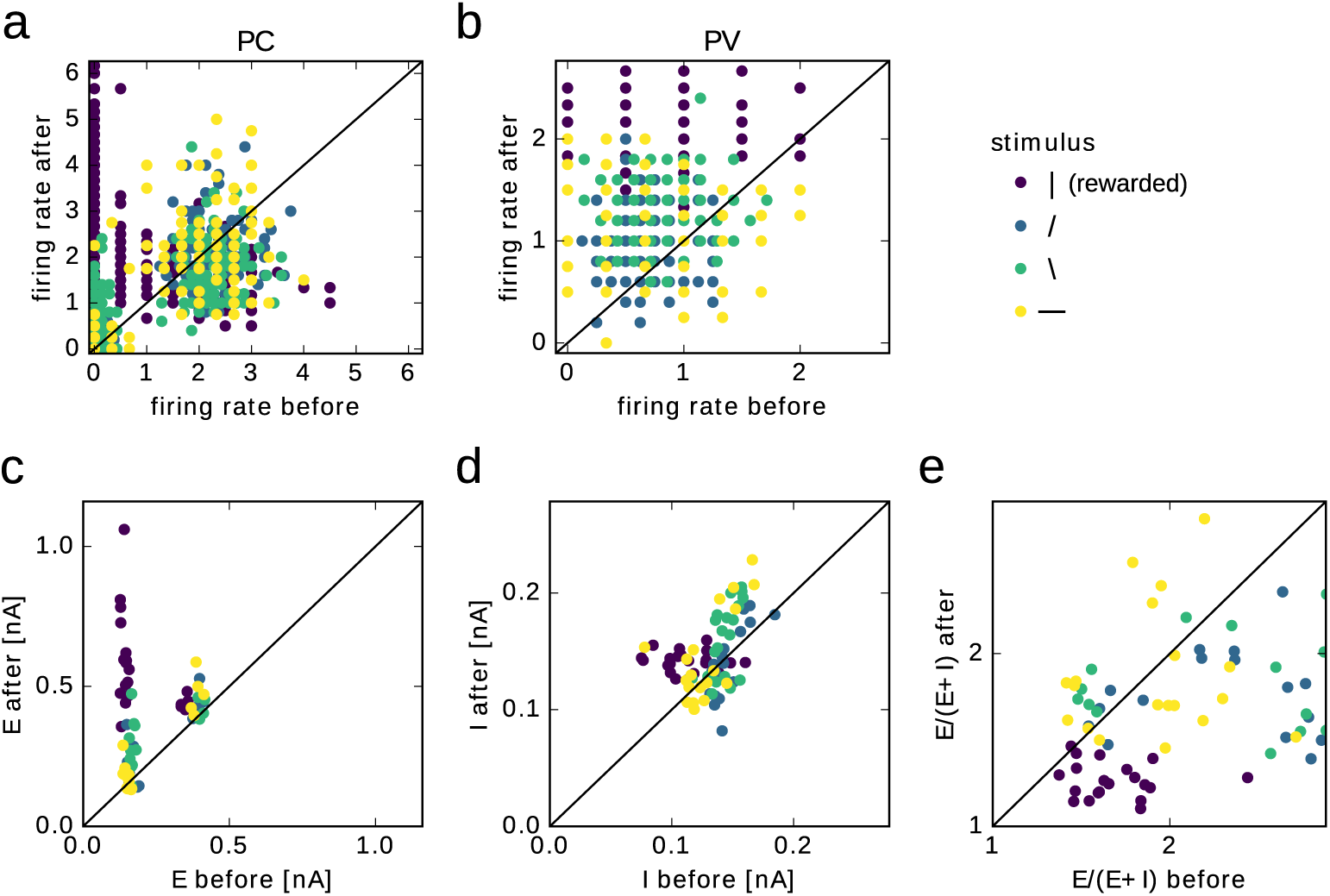
Predictions of the model. a: Firing rate of 20 sample excitatory neurons (5 from each population) before the rewarded phase as a function of after the refinement phase. b: Firing rate of sample PV interneurons. c: Excitatory currents (E), d: Inhibitory currents (I) and e: E/(E+I) ratio of the same 20 sample excitatory neurons.

### Functional implication: Translation invariance of learned representations

Most excitatory neurons in layer 2/3 are simple cells (Niell and Stryker, 2008), which respond to dedicated visual field locations. Changes in connectivity between these cells will hence only affect the representation of a stimulus at that visual field location. Therefore, we were wondering whether and how the increased representation of the rewarded stimulus could generalise to visual field locations that were not rewarded. In particular, we asked whether learning the inhibitory structure can lead to enhanced stimulus representations that are invariant to the visual field location. This so-called translation invariance is a general property of the visual system. For example, how we perceive an edge should be independent of where in the visual field it occurs.

To test this, we expanded our model to include another set of PCs, tuned to the same orientations but to a different visual location. All PCs in the model were innervated by the same set of interneurons, which were tuned to both locations (Fig. 5a) (assuming inhibition with broader spatial receptive fields, Liu et al., 2009). As before, SST-to-PV structure developed during the rewarded phase (Fig. 5e). In the refinement phase, the excitatory structure emerges in both PC subnetworks (Fig. 5b, h, and c, i), leading to an increased representation of the stimulus for both visual field locations (Fig. 5d and j). In summary, our two stage-model allows for a generalisation of the learned representation to other visual field locations.

**Figure 5:**
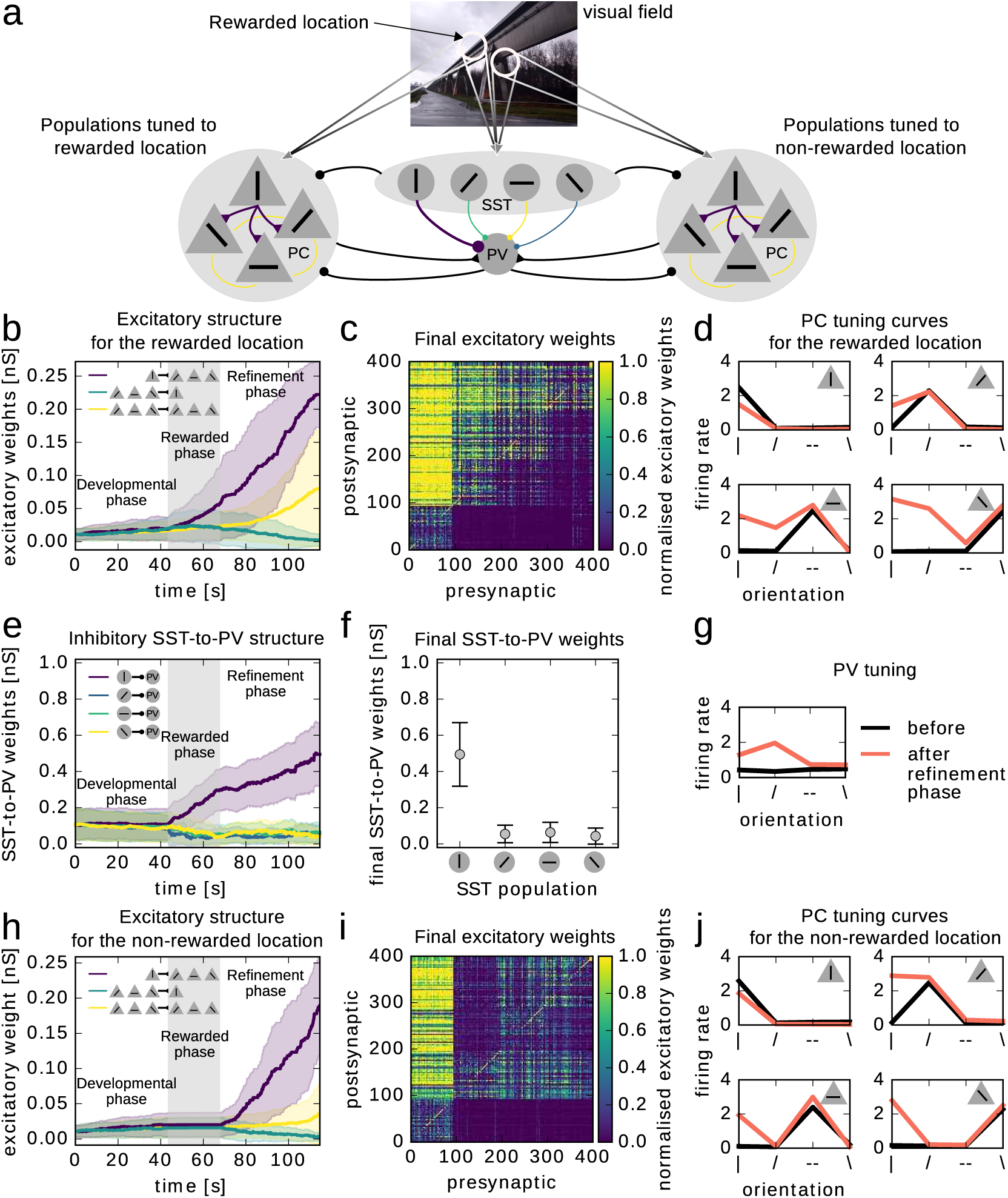
Translation invariance of learned representations. a: Illustration: Two excitatory networks with different visual receptive field locations (white circles) share the same interneuron network. The interneurons are broadly tuned, and receive input from both visual field locations. Only one visual field location is rewarded during the rewarded phase (rewarded location). b: Evolution of excitatory weights for the rewarded location. c: Final excitatory weights for the rewarded location 1. d: Tuning curves of excitatory populations with a receptive field at the rewarded location at the beginning (before) and at the end of the simulation (after the refinement phase; measured as the number of spikes during 50 ms after stimulus onset averaged over all occurrences of that stimulus in 1 s of simulation). e: Evolution of inhibitory synaptic weights. f: Final inhibitory weights. g: PV tuning at the beginning and after the refinement phase. h: Evolution of excitatory weights for the non-rewarded location. i. Final excitatory weights for the non-rewarded population. j: Tuning curves of excitatory populations with a receptive field at the non-rewarded location (as in d).

## Discussion

We propose that a memory of the rewarded stimulus is stored in the inhibitory structure. It can instruct excitatory plasticity in the absence of reward via a disinhibition mechanism. The PCs then increase their tuning to the rewarded stimulus because they receive strong connections from PCs coding for the rewarded stimulus, regardless of their initial tuning.

### Top-down signals

We show that an unspecific top-down reward signal is sufficient to create a specific circuit structure owing to the temporal coincidence between reward signals and stimulus-evoked activity. Where does the top-down signal come from? One candidate is cholinergic fibres from the forebrain, which has been shown to modulate activity in V1 (Chubykin et al., 2013; Shuler and Bear, 2006). In addition, the nucleus basalis in the basal forebrain, which sends widespread cholinergic projections to all sensory areas, has long been known to play a role in plasticity, learning, and memory. Lesioning and applying cholinergic antagonists impair learning and memory (Butt and Hodge 1995). Nucleus basalis stimulation and local ACh administration alter auditory receptive fields (Metherate and Ashe, 1991; Metherato and Weinberger, 1989). Finally, cholinergic inputs are involved in experience-dependent plasticity of visual cortex (Bear and Singer, 1986). Cholinergic inputs target many interneuron cell types (Muñoz et al., 2017). Here we focused on top-down modulation of VIPs, as VIPs (i) directly respond to reinforcement signals (Pi et al., 2013), (ii) inhibit other interneurons during learning (Fu et al., 2014; Letzkus et al., 2011), and (iii) are diversely modulated by glutamatergic, cholinergic, and serotonergic inputs (Lee et al., 2013; Prönneke et al., 2015).

### Experimental evidence demonstrating increased representation of rewarded stimuli

In Poort et al. (2015), the majority of cells increased their selectivity for one stimulus by selectively suppressing their response to the other stimuli. In Goltstein et al. (2013) on the other hand, cells increased their response to the rewarded stimulus by broadening their tuning curves. The possible discrepancy may arise from the design of the two studies. Whereas Poort et al. (2015) quantified responses to the two task-relevant stimuli, Goltstein et al. (2013) calculated tuning curves for a range of orientations including two task-relevant orientations. In our model, we captured the increased stimulus representation by an increase of responses to the rewarded stimulus, which was observed in both experimental studies (Goltstein et al., 2013; Poort et al., 2015). Additionally, PVs were also shown to increase their selectivity with learning (Khan et al., 2018).

### Limitations

Our point model does not capture the fact that SSTs and PVs target different dendritic regions. We do not expect our results to change if we include dendrites in our model, as the PV-disinhibition projects somatically. In our model, PC-to-SST connections and PV-to-PV connections were not necessary. Including them, however, yields similar qualitative results (Fig. S4), provided a homeostatic mechanism to prevent this positive feedback loop. It would be interesting to study the effect of multiple modulatory inputs, such as long-range glutamatergic, serotonergic, and cholinergic inputs. How do these inputs interact? Do they interfere with each other? How are different signals distinguished? For instance, both learning and attention affect the selectivity of responses of neurons in the circuit (Khan et al., 2018; Poort et al., 2015).

### Alternative implementations

We list below alternative mechanisms, but we argue that our model is the one most in line with experimental data from the visual cortex.

i. Vertically tuned PCs may develop strong connections to VIPs. It will inhibit SSTs and therefore disinhibit PCs. This motif, however, will cause VIPs to become more tuned during learning, which was not observed experimentally (Khan et al., 2018).
ii. Vertically tuned PCs may develop strong connections to SSTs. It will inhibit PVs and therefore disinhibit PCs. This motif will result in a tuning increase of SSTs, which was also not observed experimentally (Khan et al., 2018).
iii. Vertically-tuned PCs may reduce their inhibition of PVs, thereby increasing the activity of all PCs. This can lead to instability and contradicts the finding that PCs and PVs increase their effective connectivity during learning (Khan et al., 2018).
iv. SSTs may decrease their response to the rewarded stimulus more than to other stimuli. This motif predicts a change in the tuning of SSTs, which does not seem to be the case in Khan et al. (2018).

### Functional relevance

We propose three benefits of an intermediate inhibitory structure over direct changes in recurrent excitatory connectivity. (i) Bridging timescales: The reward is only present for a short amount of time, but plasticity can be slow. These two timescales can be bridged because the inhibitory and excitatory structure mutually reinforce each other. Therefore, a strong excitatory structure can emerge beyond the presence or even in the absence of reward. Without the inhibitory structure, the excitatory structure develops more slowly and the reward phase has to be longer (Fig. S5). Additionally, high inhibitory firing rates, typical of PVs, could effectively increase the inhibitory learning rate, allowing for a rapid development of the inhibitory structure. (ii) Translation invariance: We showed that the inhibitory structure allows for the increased representation to generalise across visual locations. Interestingly in machine learning, translation invariance which improves generalisation is achieved by weight sharing. The same weight vector (filter) is applied to different regions of the input space. This has been considered biologically implausible, as synaptic weights of far-away synapses are not locally available to each synapse. Broadly tuned interneuron networks that are shared across functional excitatory clusters may be a biologically plausible way to implement weight sharing. (iii) Stability of representations: Excitatory responses did not change during the rewarded phase. Therefore, the mechanism ensures a stable representation during relevant behaviour despite learning a structure between the inhibitory neurons.

### Conclusion

Interneuron circuits form canonical motifs across cortical areas. They integrate modulatory and long-range signals from higher cortical areas with activity in the local circuit. They are hence well-suited to adjust local circuits according to behaviourally-relevant signals. We proposed that interneuron circuits enable reward-dependent changes in sensory representations in a two-stage process. It can bridge timescales between stimulus-reward experience and synaptic plasticity. Finally, it allows for generalisation of the learned association.

## Methods

### Network model

The network consisted of 400 PCs grouped into four subpopulations of 100 neurons each. Each subpopulation coded for a given orientation. We simulated 120 PVs, 120 SST-positive interneurons (30 in each subpopulation), and 50 VIP interneurons.

### Neuron model

Neurons were modelled as conductance-based spiking leaky integrate-and-fire neurons. Their membrane potential evolves according to:

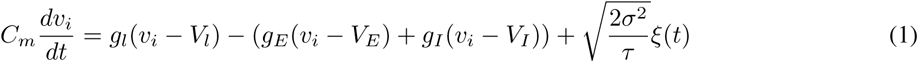

where *C*_*m*_ is the membrane capacitance, *v*_*i*_ is the membrane potential of neuron *i, V*_*l*_ is the leak reversal potential. *V*_*E*_ and *V*_*I*_ are the excitatory and inhibitory reversal potentials. *g*_*l*_, *g*_*E*_, and *g*_*I*_ are the leak, excitatory and inhibitory conductances. *g*_*E*_ and *g*_*I*_ are increased by *W*_*ij*_ upon a spike event in a pre-synaptic excitatory or inhibitory neuron *j*, and decay exponentially with time constants *τ*_*E*_ and *τ*_*I*_, respectively. ξ is zero-mean Gaussian white noise. Parameters defining the Ornstein-Uhlenbeck process are *σ* = 3 mV and correlation time *τ* = 5 ms.

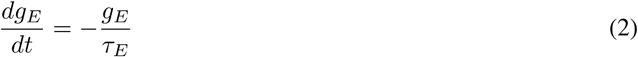

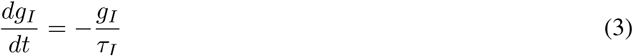

When the membrane potential reaches a threshold *v*_*θ*_, a spike event is recorded and the membrane potential is reset to its resting value *V*_*l*_.

**Table 1:** Parameters of the leaky integrate-and-fire neuron model.

| Parameter | Value | Parameter | Value |
| --- | --- | --- | --- |
| $C_m$ | 200 pF | $V_E$ | 0 mV |
| $V_l$ | -60 mV | $V_I$ | -80 mV |
| $g_l$ | 10 nS | $\tau_E$ | 5 ms |
| $v_\theta$ | -50 mV | $\tau_I$ | 10 ms |

### Connectivity

The synaptic weights *W*_*IJ*_ from a neuron *j* in population *J* to a neuron *i* in population *I*, where *I, J* ∈ {*E, P, S, V*} (*E*: PC, *P* : PV, *S*: SST, *V* :VIP), determine how much the synaptic conductances *g*_*E*_ and *g*_*I*_ increase upon a spike in neuron *j*. We initialised the synaptic weights as

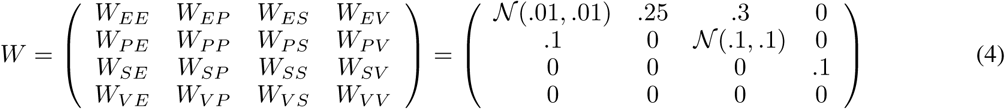

### Inputs

PCs and SSTs received one of four inputs (corresponding to layer 4 (L4) inputs coding for 4 different orientations). Each L4 input produced a Poisson-distributed spike train with a rate of 4 kHz during its preferred stimulus, and 0 Hz otherwise. One of four stimuli was shown for 50 ms followed by a stimulus gap of 20 ms. During the stimulus gap, all L4 inputs produced spikes at the same rate of 1.6 kHz. The conductance of synapses from L4 to PCs was .25 nS. The conductances of synapses from L4 to SSTs was .2 nS during the stimulus and .22 nS during the stimulus gap. Additionally, PCs and PVs received a baseline input from a Poisson process with a rate of 4 kHz. The weights to PCs weights were .06 nS and to PVs were .01 nS.

### Plasticity

For both excitatory and inhibitory plasticity we chose the simple classical STDP model (Bi and Poo, 1998; Gerstner et al., 1996; Markram et al., 1997)

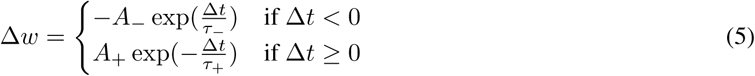

where Δ*t* = *t*_post_ - t_pre_ is the difference between pre- and postsynaptic spike time, *τ*_+_ = *τ* = 20 ms, for excitatory plasticity *A*_+_ = .005 nS and *A* =1.05*A*_+_, for inhibitory plasticity *A*_+_ = .015 nS and *A* =1.05*A*_+_. This rule leads to synaptic potentiation when the presynaptic neuron spikes before or simultaneously with the postsynaptic neuron, and to depression otherwise.

In the online implementation of this rule, the synaptic weight *w*_*ij*_ from neuron *j* to neuron *i* is updated when either the pre- or the postsynaptic neuron spikes according to:

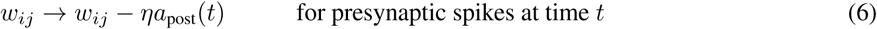

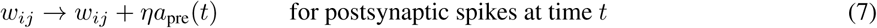

where *a*_post_(*t*) is the postsynaptic trace and *a*_pre_(*t*) is the presynaptic trace. The traces are updated by a constant value at the time of a postsynaptic or presynaptic spike respectively, and decay exponentially with a time constant *τ*_*-*_ or *τ*_+_.

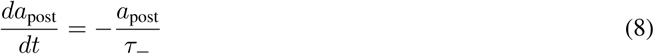

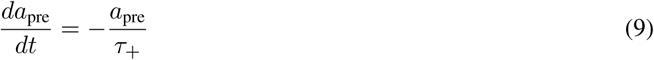

Excitatory and inhibitory weights were constrained to be positive and had an upper bound at 0.25 nS for excitatory weights and 1 nS for inhibitory weights.

### Simulation

All spiking simulations were done with the Brian 2 simulator (Stimberg et al., 2013), using a time step of 0.1 ms. The model was simulated for 1.4 s without plasticity to measure tuning curves. Then plasticity was switched on. The model was simulated for 42 s during the developmental phase, followed by a 24.5 s during the rewarded phase, and 45.5 s during the refinement phase. Finally, the model was simulated for 1.4 s without plasticity to measure final tuning curves again.

### Translation invariance

To adjust for the increased number of excitatory neurons in the network, the connection strength from PCs to PVs was decreased to *W*_*P E*_ = .06 nS.

### Data availability

All simulation code used for this paper will be made available on ModelDB.

## Supplementary Material

### Spiking information

To understand the sources of the synaptic changes, we analysed the spike timing of the different neuron types. (i) During the rewarded phase, VIPs were excited when the vertical bar was shown. As VIPs suppress SSTs, SSTs fired only briefly to the stimulus before they were silenced by VIPs. The lack of SST inhibition caused an increase in PV firing after the SSTs fired (Fig. S1b middle), such that spikes in SSTs preceded spikes in PVs during the vertical bar (Fig. S1d vertical SST - PV). As a consequence, connections from vertically tuned SSTs to PVs increased (Fig. 2b). (ii) During the refinement phase, the strong vertically tuned SST to PV connections led to increased inhibition of PVs. This disinhibited the PCs when a vertical bar was shown. As a result of this disinhibition, they increased their response towards the vertical bar and their likelihood to fire together. Additionally, vertically tuned PCs fired before the others as they received additional feedforward sensory input (Fig. S1c top and Fig. S1e vertical PC - angled PC). Therefore, vertically tuned PCs to other PCs connection strengthened. At the same time, the increased firing of PCs to the vertical bar increased PV firing just after vertically tuned SSTs responded. The vertically tuned SST-to-PV connections hence further strengthened (Fig. 3f).

**Figure S1:**
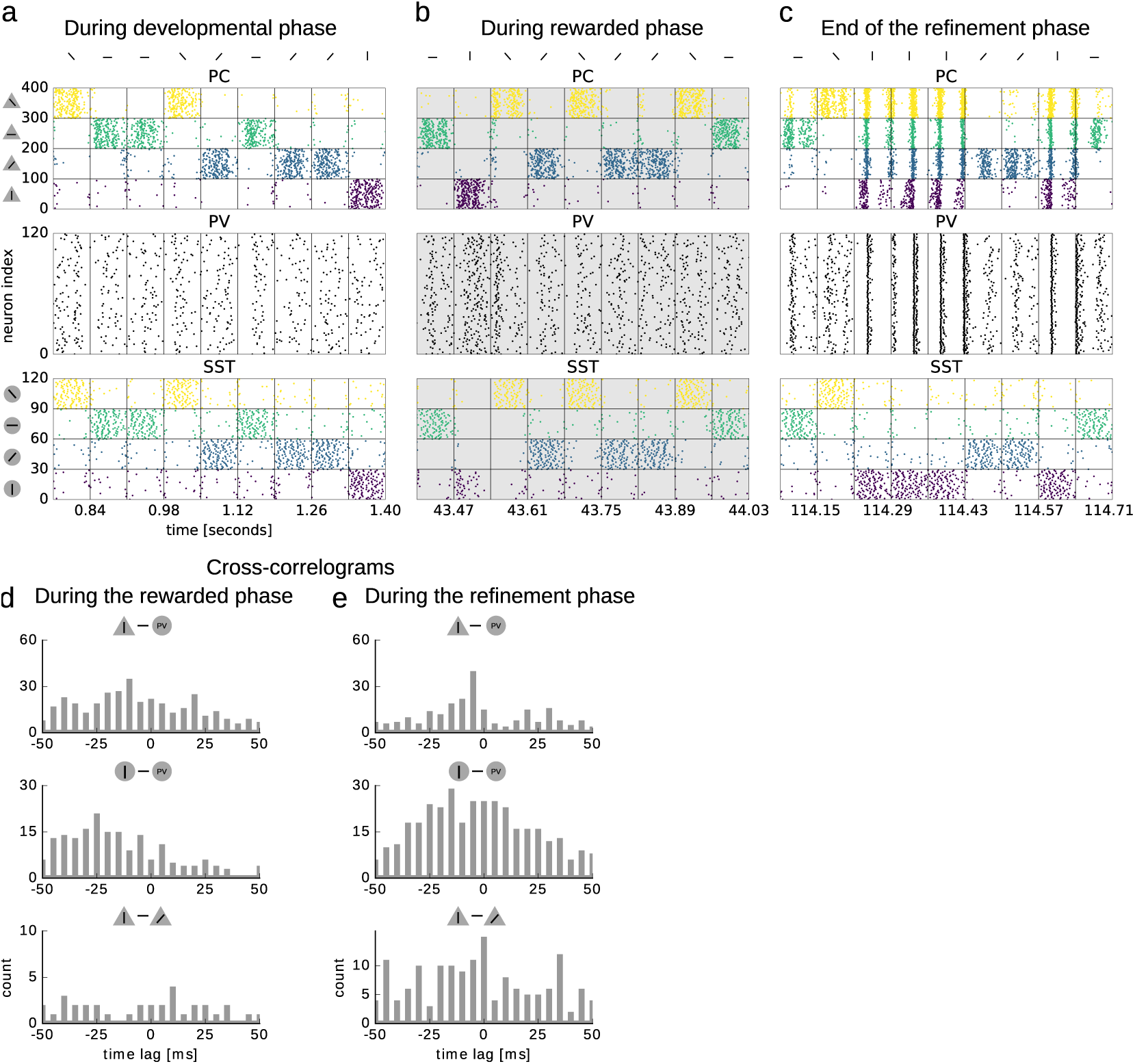
Spiking information. a-c: Spike raster plots of PCs (top), PV interneurons (middle) and SSTs (bottom) during different stages of the simulation: at the beginning of the rewarded phase (a), during the rewarded phase (b, grey background), and in the end of the rewarded phase (c). Sensory inputs are presented for 50 ms, followed by a 20 ms stimulus gap. Changes of stimulus are indicated by vertical lines. d,e: Cross-correlograms for pairs of cells from different cell classes during (d) and after (e) the rewarded phase. Top: vertically tuned PC and PV. Middle: vertically tuned SST and PV. Bottom: vertically tuned PC and angled tuned PC.

### SST tuning properties do not change

**Figure S2:**
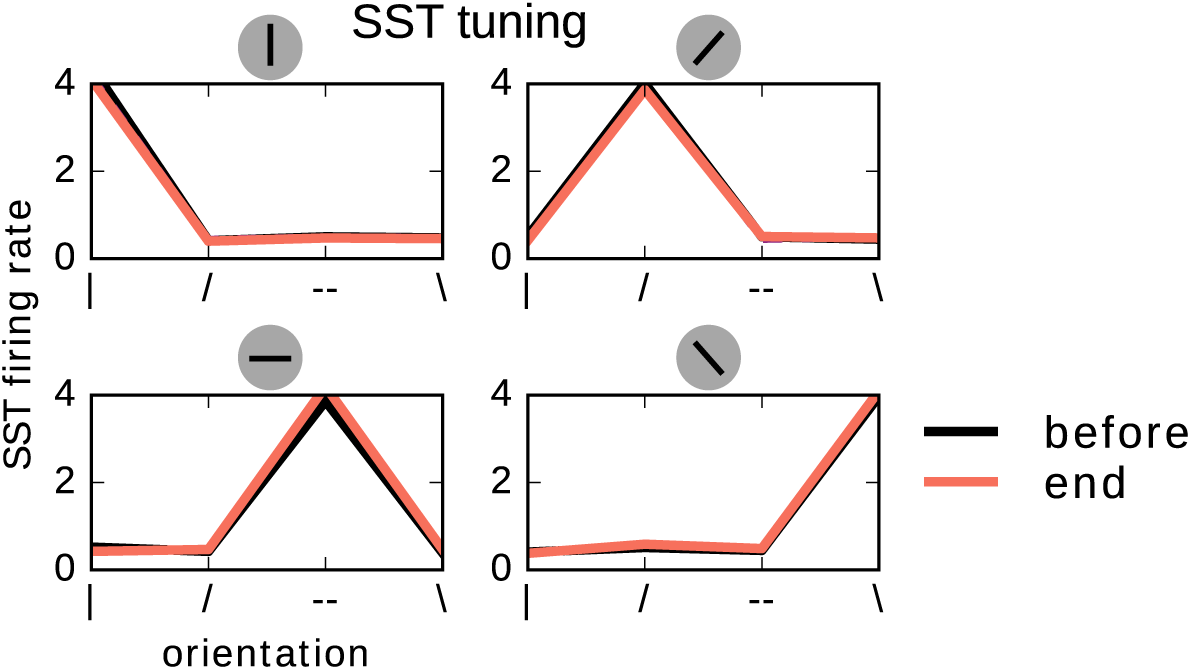
SST tuning in spiking model do not change. Tuning of SST populations before the rewarded phase (black) and at the end of the refinement phase (red, number of spikes during 50 ms after stimulus onset averaged over all occurrences of that stimulus in 1 s of simulation). Tuning preference of each population is indicated above each panel.

### The two-stage model is robust across several implementations

**Figure S3:**
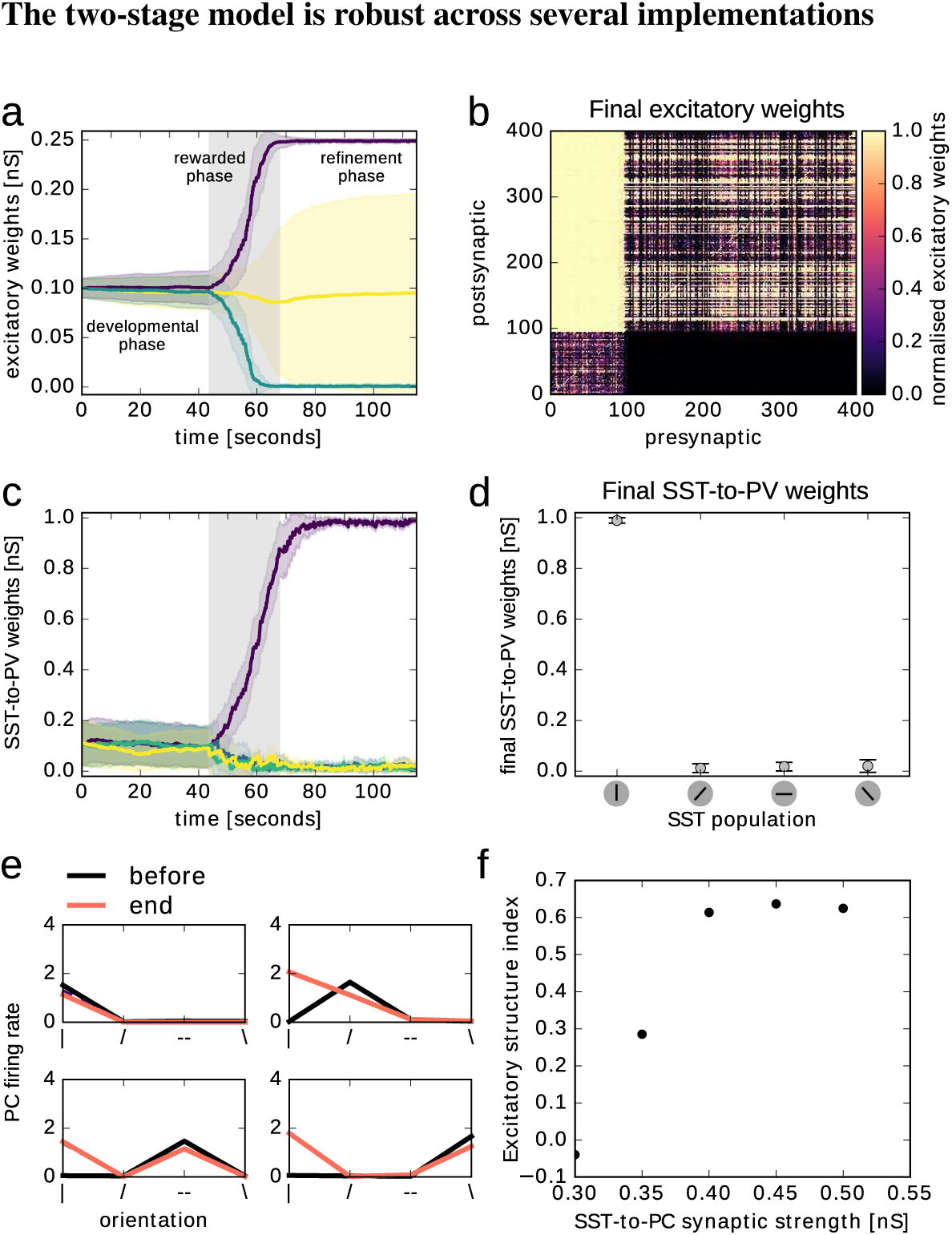
Spiking model with strong initial recurrent connectivity. Inhibitory and excitatory structure also developed with strong recurrence: *W*_*EE*_ = 𝒩(*µ* = .1, *σ* = .01). SST-to-PC connections needed to be strong to keep the network stable: W_*P C,SST*_ = .5 nS a: Evolution of excitatory connections (mean and standard deviation). Non-vertically tuned PCs to vertically tuned PCs (green), vertically tuned PCs to non-vertically tuned PCs (purple), non-vertically tuned PCs to non-vertically tuned PCs (yellow). b: Recurrent excitatory weights at the end of the refinement phase. c: Evolution of the inhibitory SST-to-PV connections, grouped according to SST tuning preference (colours as in Fig.2b, vertical in purple), shown are the mean and standard deviation. d: SST-to-PV weights (mean and standard deviation) at the end of the refinement phase averaged over SST tuning preference. e: Tuning of excitatory populations before the rewarded phase (black) and at the end of the refinement phase (red, number of spikes during 50 ms after stimulus onset averaged over all occurrences of that stimulus in 1 s of simulation). f: Excitatory structure index (see methods) as a function of SST-to-PC synaptic strength.

**Figure S4:**
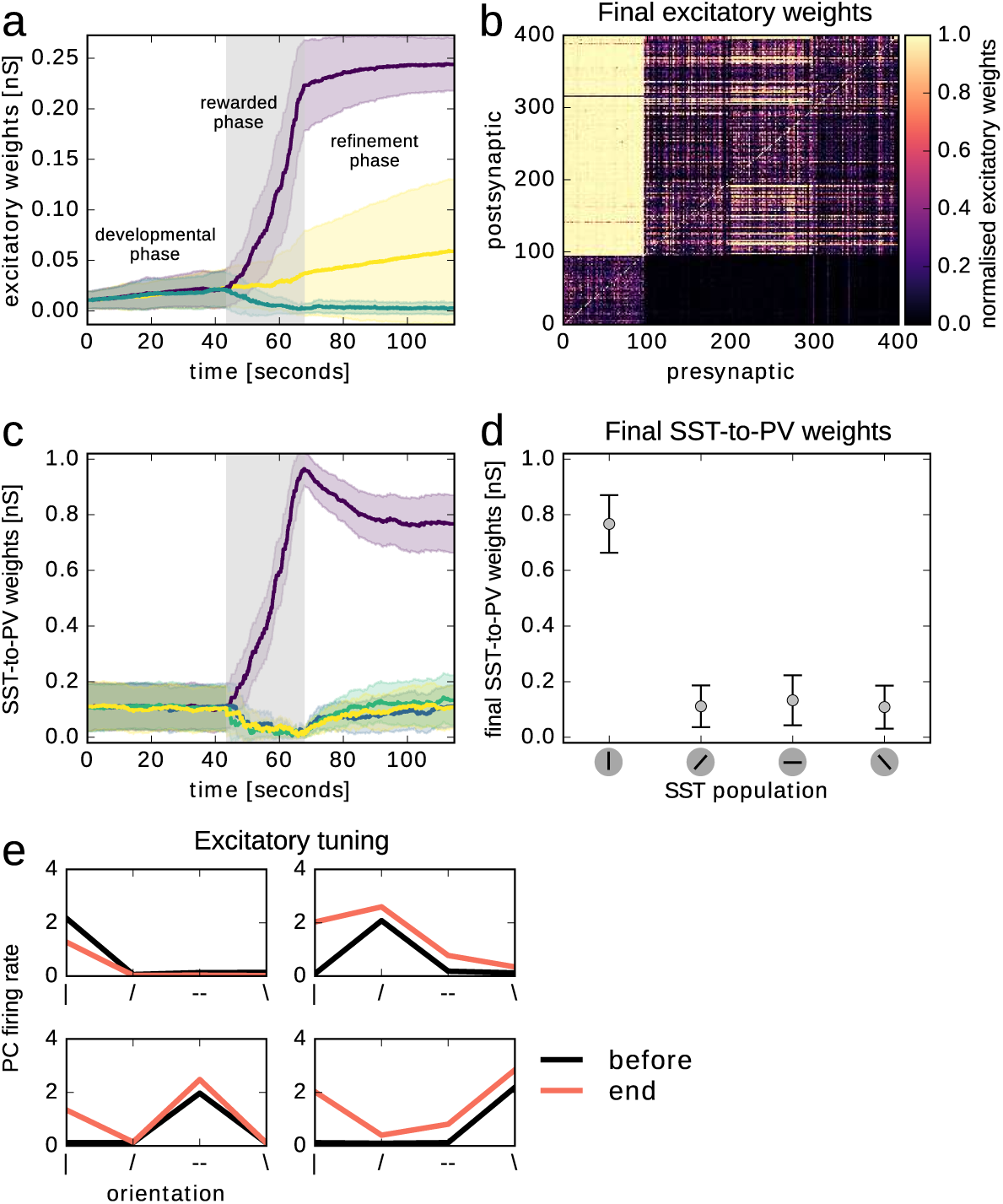
Spiking model with PC-to-SST and PV-to-PV connections. Introducing PC-to-SST connections can lead to a strong disinhibition via the SST-PV-PC pathway, which can be counteracted by PV-to-PV connectivity. PV-to-PV self-inhibition effectively weakens the SST-PV-PC pathway. The relative increase in SST-PC strength (and hence stronger disinhibition during the reward) is reflected by the development of excitatory connectivity during the rewarded phase. Instabilities introduced by recurrent loops (here PC-SST-PV-PC) can alternatively be accounted for by homeostatic mechanisms that regulate connectivity strengths to keep the network in its operating mode. W_*SST,P C*_ = .13 nS and W_*PV,PV*_ = .06 nS. a: Evolution of excitatory connections (mean and standard deviation). Non-vertically tuned PCs to vertically tuned PCs (green), vertically tuned PCs to non-vertically tuned PCs (purple), non-vertically tuned PCs to non-vertically tuned PCs (yellow). b: Recurrent excitatory weights at the end of the refinement phase. c: Evolution of the inhibitory SST-to-PV connections, grouped according to SST tuning preference, shown are the mean and standard deviation. d: SST-to-PV weights (mean and standard deviation) at the end of the refinement phase averaged over SST tuning preference (colours as in Fig.2b, vertical in purple). e: Tuning of excitatory populations before the rewarded phase (black) and at the end of the refinement phase (red, number of spikes during 50 ms after stimulus onset averaged over all occurrences of that stimulus in 1 s of simulation).

### With the development of inhibitory structure, shorter reward duration is needed for the development of excitatory structure

**Figure S5:**
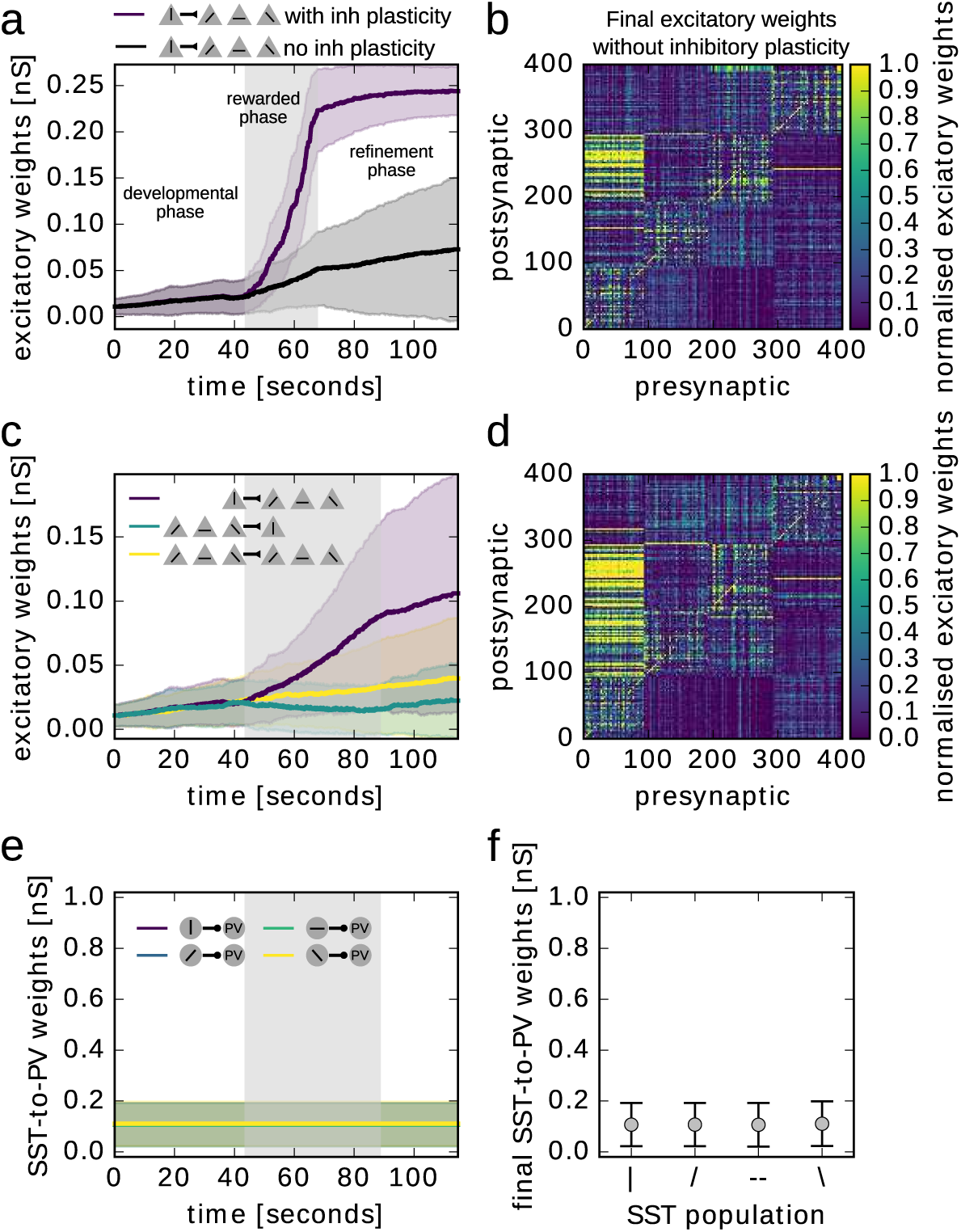
Spiking model without inhibitory plasticity. Excitatory structure develops more slowly without inhibitory plasticity. For a fair comparison, we used the model in which excitatory structure already develops during the rewarded phase shown in Fig. S4 with W_*SST,P C*_ = .13 nS and W_*PV,PV*_ = .06 nS. a: Evolution of excitatory connections (mean and standard deviation). Development of connections from vertically tuned PCs to non-vertically tuned PCs in the model without inhibitory plasticity (black) compared to the model with inhibitory plasticity (purple). b: Recurrent excitatory weights at the end of the refinement phase in the model without inhibitory plasticity. c-f: Model without inhibitory plasticity and a longer reward phase. c: Evolution of excitatory connections (mean and standard deviation) in the model without inhibitory plasticity: non-vertically tuned PCs to vertically tuned PCs (green), vertically tuned PCs to non-vertically tuned PCs (purple), non-vertically tuned PCs to non-vertically tuned PCs (yellow). d: Recurrent excitatory weights at the end of the refinement phase in the model without inhibitory plasticity. e: Evolution of the inhibitory SST-to-PV connections, grouped according to SST tuning preference, shown are the mean and standard deviation. f: SST-to-PV weights (mean and standard deviation) at the end of the refinement phase averaged over SST tuning preference (colours as in Fig.2b, vertical in purple).

### Our two-stage model is robust to perturbations: excitatory structure remains although the inhibitory structure is artificially reset

**Figure S6:**
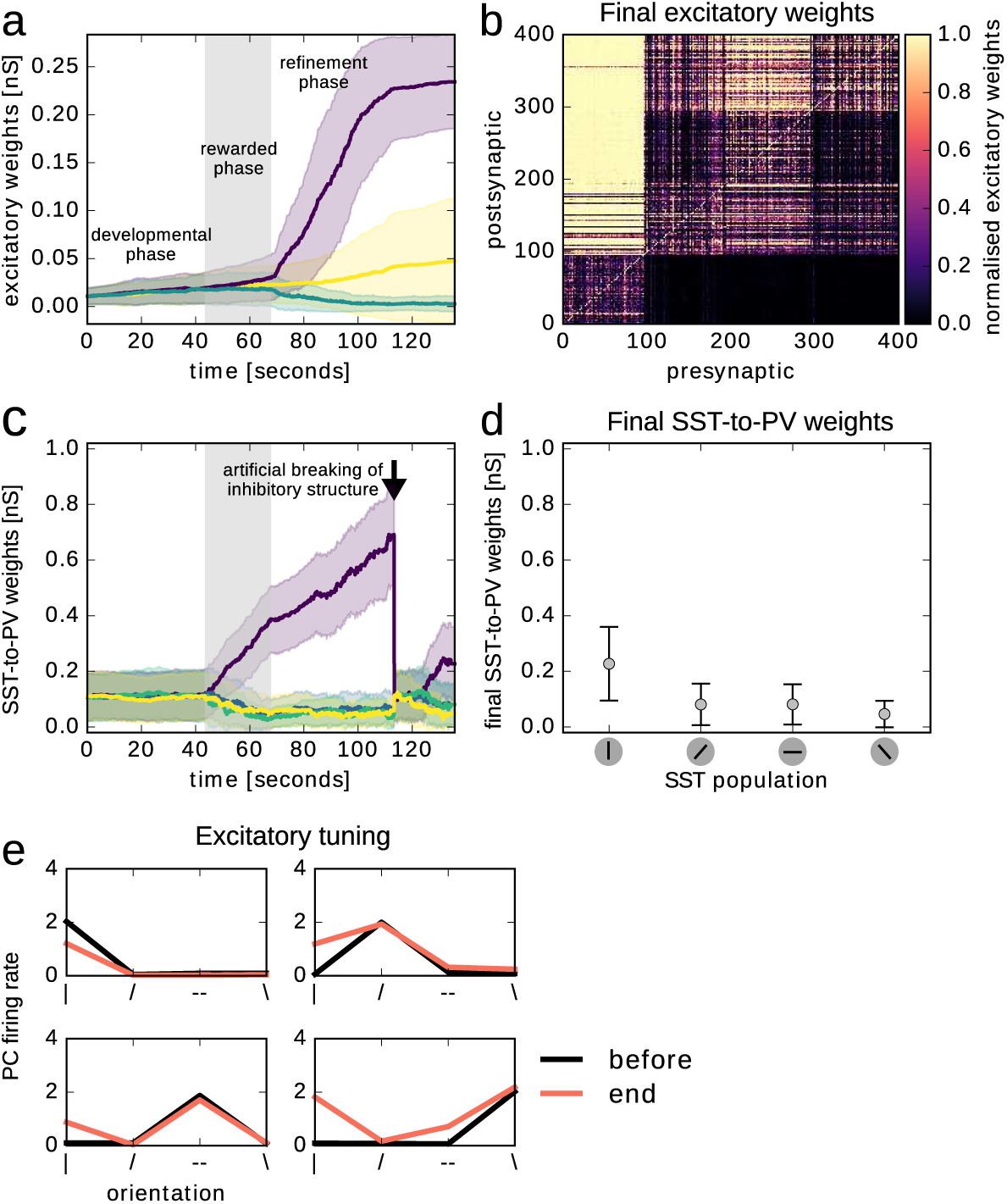
Excitatory structure is stable without inhibitory structure. Simulation with same parameters as Fig. 2, but SST-to-PV connections were reset to their initial values at time 113.4 s. The excitatory structure remained stable. a: Evolution of excitatory connections (mean and standard deviation). Non-vertically tuned PCs to vertically tuned PCs (green), vertically tuned PCs to non-vertically tuned PCs (purple), non-vertically tuned PCs to non-vertically tuned PCs (yellow). b: Recurrent excitatory weights at the end of the refinement phase. c: Evolution of the inhibitory SST-to-PV connections, grouped according to SST tuning preference (colours as in Fig.2b, vertical in purple), shown are the mean and standard deviation. d: SST-to-PV weights (mean and standard deviation) at the end of the refinement phase averaged over SST tuning preference. e: Tuning of excitatory populations before the rewarded phase (black) and at the end of the refinement phase (red) (number of spikes during 50 ms after stimulus onset averaged over all occurrences of that stimulus in 1 s of simulation).

### Precise spike timing is not necessary: The two-stage model can be implemented in a rate coding scheme

Although precise spike timing (Tiesinga et al., 2008) and spike-timing based plasticity rules (Sjöström et al., 2001) have been reported in the primary visual cortex, it is unknown whether they play a role in enhancing stimulus representations. We hence investigated whether our two-stage model can also be implemented in a rate-based framework. Our network was in the inhibition-stabilized regime (ISN, Tsodyks et al. (1997), recent experimental support: Moore et al. (2018)) and the learning rule was a BCM type. In this regime, PC firing ‘paradoxically’ increases with increased PV activity. This is due to strong recurrent excitatory connections balanced by strong inhibition. Suppression of the inhibitory population in an ISN leads to an immediate increase in excitatory activity, which in turn drives the inhibitory population.

We implemented excitatory and inhibitory rate-based plasticity rules such that weight changes resembled those in the spiking implementation. (i) Excitatory plasticity favoured connections from high firing neurons to low firing neurons and depressed the reverse (see supplementary methods). The SST-to-PV plasticity potentiated synapses active while PVs fired above a threshold 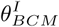, reminiscent of the BCM learning rule (Bienenstock et al., 1982).

During the rewarded phase, the SST-to-PV connectivity developed akin to the spiking model. In particular, connections from the vertically tuned SSTs to PVs strengthened, while the other connections weakened (Fig. S7b bottom, grey background). Additionally, the connection from the non-rewarded PCs to the rewarded vertically tuned PCs decreased (green line in Fig. S7b top).

During the refinement phase, PVs received more inhibition during the vertical stimulus (due to the strong connection from the vertically tuned SSTs). Nevertheless, PV firing rate was higher during the vertical bar than during the horizontal stimulus (Fig. S7f around reward end). This reflects the ISN property of the network. Hence, the connection from the vertically tuned SST population continued to increase and remained stronger (Fig. S7b bottom) than the connection from the other SSTs. With the inhibitory structure in place, the vertically tuned PCs to horizontally tuned PCs connection strengthened (purple line in Fig. S7b top). Hence, the inhibitory structure remained stable and excitatory structure developed during the refinement phase. The resulting excitatory connectivity resembles that in the spiking model (Fig. S7c).

As in the spiking model, both PCs and the PVs became more tuned to the rewarded stimulus (Fig. S7d and e). The excitatory populations simply increased their firing rate towards the rewarded stimulus (Fig. S7d). The PVs, however, increased their tuning by firing less towards the non-rewarded stimulus (Fig. S7e). The decrease in PV firing towards the non-rewarded stimulus resulted from an increased SST-to-PV connectivity from both SST populations.

Even though the spiking implementation yields similar results for weak and strong initial connectivity (Fig. S3), the rate implementation requires strong recurrent connectivity. To conclude, the proposed two-stage model can be realised even if only rate information is available. This happens provided that the network exhibits a counterintuitive increase of inhibitory firing rates, a property of inhibition-stabilised networks (Tsodyks et al., 1997).

In the spiking model, inhibitory currents increase during the rewarded stimulus (Fig. 4d), whereas in the rate model inhibitory currents decrease during the non-rewarded stimulus (Fig. S7h). To delineate the spiking implementation from the rate coding implementation, the spiking model makes further predictions: (i) PVs fire more in synchrony (Fig. S1c middle), (ii) SSTs fire before PVs during the rewarded stimulus (Fig. S1d), (iii) PCs tuned to the rewarded stimulus fire before others during the rewarded stimulus after the task (Fig. S1e).

**Figure S7:**
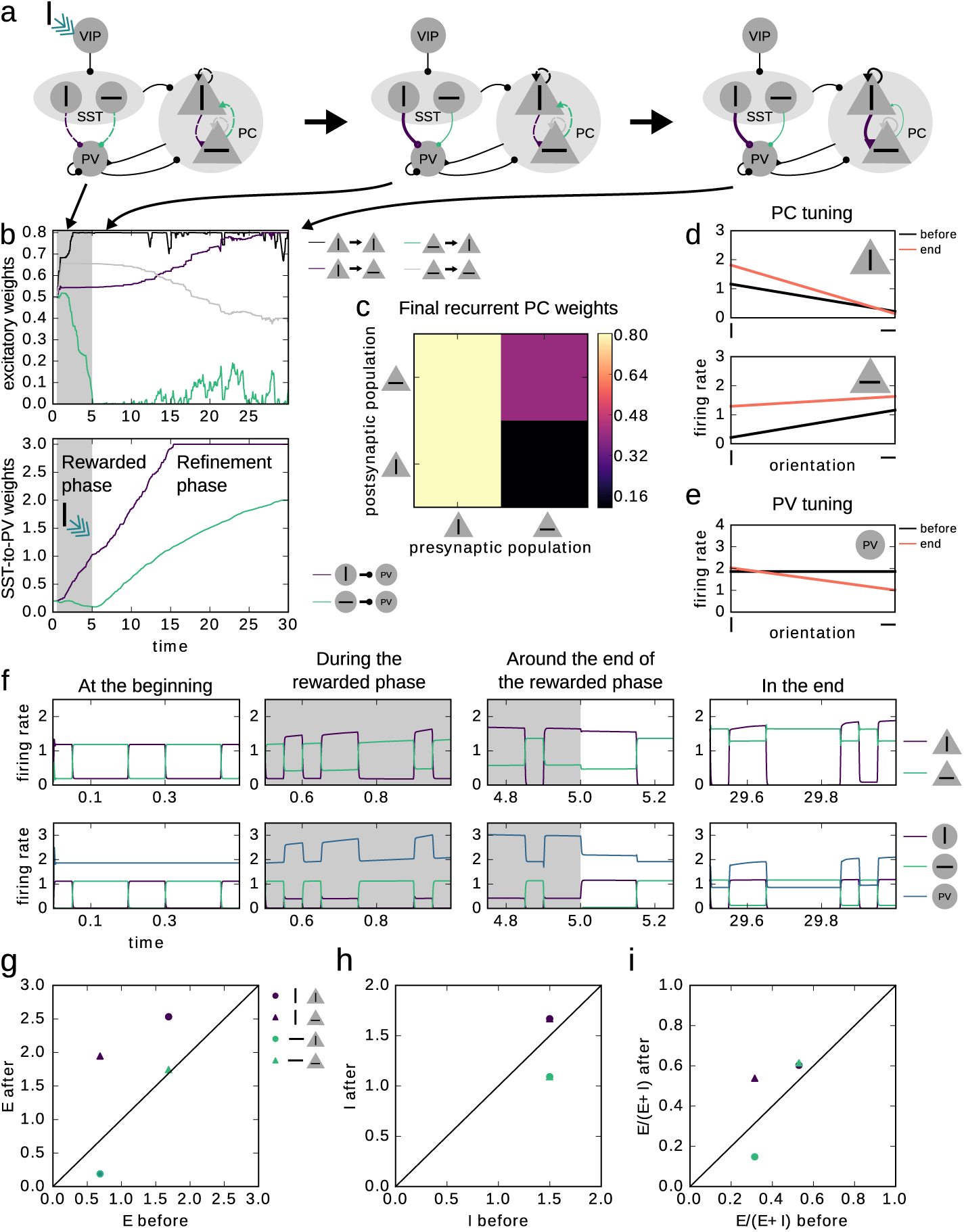
The two-stage model of microcircuit plasticity can be realised in a rate coding scheme. a: Illustration of changes in inhibitory and excitatory connectivity. Snapshots of connectivity at the beginning (left), after the rewarded phase (middle) and at the end of the refinement phase (right). b: Top: Evolution of excitatory connections from the vertically tuned PC population to itself and to the horizontally tuned PCs, and from horizontally tuned PCs to horizontally tuned PCs (grey) and to vertically tuned PCs (green). Bottom: Evolution of connections from the vertically tuned SSTs to PVs (purple) and from the horizontally tuned SSTs to PV (green). Grey background highlights rewarded phase. C: Final excitatory connectivity matrix. d: Tuning curves of excitatory populations at the beginning and at the end of the simulation. e: Tuning curves of PV population at the same time points as in d. f: Firing rates of excitatory (top) and inhibitory (bottom) populations at the beginning of the simulations, during the rewarded phase, towards the end of the rewarded phase, and at the end of the simulation (from left to right). g-i: Excitatory (g), inhibitory (i), and E/(E+I) ratio (i) in excitatory populations (circles: vertically tuned, triangles: horizontally tuned) of rate model during the two different stimuli (purple: vertical stimulus, green: horizontal stimulus) before and after learning.

## Supplementary Method

### Excitatory structure index

The excitatory structure index is defined as the difference between the mean weight from vertically tuned PCs to all other neurons and the mean weight from PCs not coding for vertical bars to vertically tuned PCs, normalised by the maximum weight:

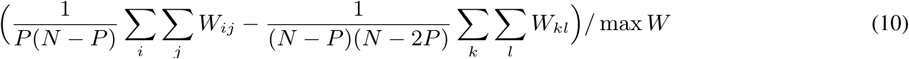

with j indexing vertically tuned PCs (1 *< j < P*), and i indexing other PCs (*i > P*). If *k* ∈ *K* then *l* ∉ *K*, and *k, l > P*, where P is the number of PCs in one population, and N is the total number of PCs.

### Rate model

The activity of a population of neurons *i* is described by their firing rate *r*_*i*_, which evolves over time according to

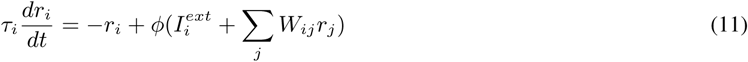

where *i, j ∈* {*E*1, *E*2, *P, S*1, *S*2, *V*} and *τ*_*i*_ is the time constant of population *i*. 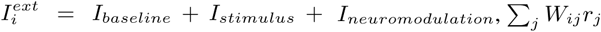 is recurrent input. φ(*x*) is the activation function given by

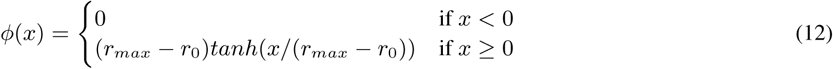

Each of the PCs and SSTs receive an input current *I*_*stimulus*_ upon presentation of their preferred stimulus. PCs also receive a constant baseline current input *I*_*baseline*_. The VIPs receive a neuromodulatory current *I*_*neuromodulation*_ when the rewarded stimulus (vertical bar) is present. *r*_0_ and *r*_*max*_ denote the minimum and maximum firing rate, respectively.

### Connectivity

The initial connectivity is taken such that the network is in the ISN regime.

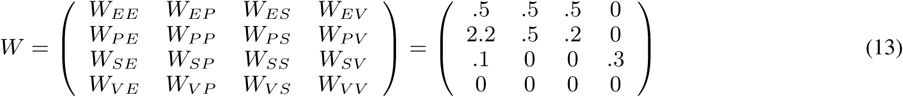

### Plasticity rules

#### Excitatory Plasticity

For excitatory connections, we used a BCM-like rule (Bienenstock et al.1982), where the sign of synaptic change depends on whether the activity of the postsynaptic neuron exceeds a threshold. The rule is Hebbian as the weight change depends on the product of pre- and post-synaptic activity. The weight change follows

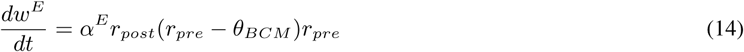

where *α*^*E*^ is the excitatory learning rate, *r*_*pre*_ is the presynaptic firing rate, *r*_*post*_ the postsynaptic firing rate, and 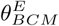 is the sliding threshold. The threshold is sliding and changes according to:

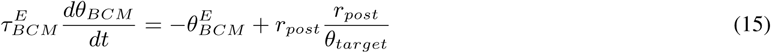

where *θ*_*target*_ is the target firing rate and 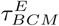 the time constant.

#### Inhibitory Plasticity

For inhibitory connections, we used:

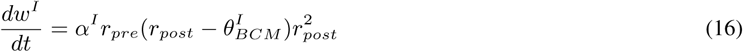

where *α*^*I*^ is the learning rate, *r*_*pre*_ is the presynaptic firing rate, *r*_*post*_ the postsynaptic firing rate, and 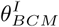 is the inhibitory sliding threshold. It changes according to:

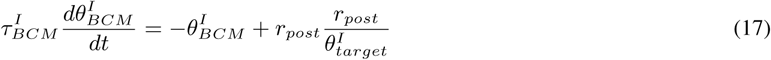

where *θ*_*target*_ is the target firing rate and 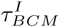 the time constant.

**Table S1:** Parameters of the rate model.

| **Parameter** | **Value** | **Parameter** | **Value** |
| --- | --- | --- | --- |
| $\tau_i$ | 1.0 | $\alpha^E$ | 5.0e-4 |
| $r_0$ | 1.0 | $\tau_{BCM}^E$ | 1/1.0e-2 |
| $r_{\max}$ | 20.0 | $\theta_{\text{target}}$ | 1.5 |
| $I_{\text{stimulus}}$ | 1.0 | $W_{\text{maxsum}}^I$ | 5.0 |
| $I_{\text{baseline}}$ | 1.0 | $W_{\text{max}}^I$ | 3.0 |
| $I_{\text{neuromodulation}}$ | 2.5 | $\alpha^I$ | 1.0e-4 |
| $W_{\text{max}}$ | .8 | $\tau_{BCM}^I$ | 1/1.0e-3 |
| $W_{\text{maxsum}}$ | 1.2 | $\theta_{\text{target}}^I$ | 3.2 |

### Simulation

All rate model simulations were done with a time step of 0.1 [arb. unit]. First, the model was simulated for 500 [arb. unit] without plasticity to measure the tuning properties. Then plasticity was switched on, and the rewarded phase started. The rewarded phase ended at time 5000 [arb. unit]. The refinement phase ended at time 30000 [arb. unit].

